# Biocompatible fluorocarbon liquid underlays for in situ extraction of isoprenoids from microbial cultures

**DOI:** 10.1101/2022.01.27.477974

**Authors:** Sebastian Overmans, Kyle J. Lauersen

## Abstract

Microbial production of heterologous metabolites is now a mature technology in many host organisms, opening new avenues for green production processes for specialty chemicals. At lab scale, petroleum-based hydrophobic bio-compatible solvents like dodecane can be used as a second phase on top of microbial cultures to act as a physical sink for heterologous hydrocarbon products like isoprenoids. However, this approach has significant drawbacks at scale due to the difficulty of handling solvents and their potential contamination with unwanted byproducts of their manufacture. We discovered that synthetic perfluorocarbon liquids (FCs), commonly used for heat transfer, can also act as physical sinks for microbially produced isoprenoid compounds. FCs are stable, inert, and are amenable to direct liquid-liquid extraction with alcohols for rapid product isolation. These liquids are more dense than water and form a lower phase to microbial cultures rather than an upper phase as with other solvents. Their ability to form an under-layer or ‘underlay’ also enables the cultivation of microbes directly at the FC-culture medium interface via gravity settling, which could open their application for filamentous or mat-forming organisms. We present comparisons of the isoprenoid extraction potential of three commercial FCs: FC-3283, FC-40, and FC-770 with engineered green microalga cultures producing patchoulol, taxadiene, casbene, or 13R(+) manoyl oxide. We demonstrate that FCs are promising alternatives to traditional solvents and open new avenues in bio-process design for microbial heterologous metabolite milking.

## Introduction

Metabolic engineering of microbes for the production of heterologous isoprenoids is now a mature technology, with several commercial enterprises converting base feedstocks into hydrocarbons of various complexities (Daletos et al., 2020). Hemi- (C5), mono- (C10), sesqui- (C15), di- (C20), tri- (C30), non-canonical terpenoids (C11, C12, C16, C17), and carotenoids (Heider et al., 2012) are now routinely produced in diverse host organisms such as bacteria, yeasts, microalgae, and mosses (Lauersen, 2019; Navale et al., 2021; Wolf et al., 2021; Zhan et al., 2014). These engineering successes leverage the universal need for isoprenoid precursors isopentenyl and dimethylallyl diphosphate (IPP and DMAPP), and their subsequent condensed and dephosphorylated hydrocarbon chains. Core isoprenoids have cellular roles in membrane fluidity, sterol biosynthesis, prenylation, light capture, photoprotection, and as electron carriers (Wichmann et al., 2020). The base hydrocarbon backbones of isoprenoids have also served as a substrate for the evolution of complex specialty chemical structures through the evolution of terpene synthases that mediate deprenylation, cyclization, folding, and (sometimes) hydroxylation (Gershenzon and Dudareva, 2007; Kirby and Keasling, 2009). Base terpenes can be further chemically decorated by the action of cytochrome P450s, which add functional groups to increase their chemical diversity (Jensen et al., 2021; Wlodarczyk et al., 2016). Terpene synthases have evolved in bacteria (Yamada et al., 2015), fungi (Wei et al., 2020), plants (Kirby and Keasling, 2009), insects (Beran et al., 2016; Gilg et al., 2009), and some algae (Wei et al., 2020) as a means to generate specialty chemicals from base isoprenoids IPP and DMAPP to be used in signaling, defense, and attraction/repulsion.

Many isoprenoids are naturally produced in small quantities and may be found on slow-growing, non-farmable, or environmentally sensitive organisms (Gershenzon and Dudareva, 2007). The biological universality of isoprenoid precursors IPP and DMAPP enables the transfer of modular terpene pathways from progenitors to microbial hosts by the heterologous expression of their terpene synthases and P450s (Kirby and Keasling, 2009). Engineered microbes serve this role well and can be used to scale the production of isoprenoid(s) from their biomass in fermenters and bioreactors (Li et al., 2020). At lab-scale, microbial isoprenoid productivity can be quantified using hydrophobic biocompatible solvent overlays as a two-phase constant extraction method on living cultures. These solvents act as a physical sink to simultaneously remove and capture heterologous isoprenoids from microbial cells; a process sometimes referred to as ‘milking’. Some commonly used solvents include decane (Peralta-Yahya et al., 2011), dodecane (Beekwilder et al., 2014), or isopropyl myristate (Liu et al., 2016; Yunus and Jones, 2018). These act similarly to the trichomes of plants by accumulating the hydrophobic isoprenoid products, alleviating product inhibition, and promoting forward reactions while serving as a simple-to-separate phase for easy product analysis.

Hydrocarbon solvent-medium dual-phase concepts are convenient at small scales, however, they have significant drawbacks that prevent their use at volumes greater than a few hundred milliliters. Alkane solvents are flammable and require specialized infrastructure for pumping, as they are incompatible with silicone-based tubing. These solvents are also distillates of petroleum, which can contain many contaminants depending on the supplier. The most notable issue with using hydrophobic solvent overlays is the difficulty in scaling their application to larger culture volumes in turbid and sparged bioreactor concepts (Lauersen, 2019). As an overlay, any turbidity at the culture-solvent interface creates an emulsion of hydrophobic cellular components and the solvent that can blow out of reactors or lyse cells, hindering the processibility and productivity of extracted isoprenoids.

The lack of translatability of these lab processes to scale leaves significant room for improvement. Here, we show that long-chain hydrophobic perfluorocarbons that are liquid in operational temperature ranges can act as alternatives to hydrocarbon solvents for microbial culture milking. These liquids have current application as heat-transfer fluids and are used in cooling systems for electronics (Choi and Cho, 2000; Jouhara and Robinson, 2010). As they are more dense than water, perfluorocarbon liquids settle as a lower-phase or ‘underlay’ below the culture. Perfluorocarbon liquids are inert, available in a range of chemical conformations with high heat stability, and significantly easier to handle than hydrocarbon solvents. We show that perfluorocarbon liquids provide cleaner extraction of isoprenoid products from microbial culture than dodecane, fluorocarbon type can be tailored to target isoprenoid properties, and their extreme densities enable subsequent liquid-liquid extraction of collected terpenoids with alcohols. Hydrophobic underlays also provide unique opportunities for the direct cultivation and milking of biofilms at the medium-fluorocarbon interface.

## Materials and Methods

### Chemicals

Three perfluoro liquids, hereafter referred to as FCs, were used in this work: The perfluorinated amine Fluorinert™ FC-40 was purchased from Sigma-Aldrich (Overijse, Belgium), while the perfluorinated ether FC-770 and the perfluorinated amine FC-3283 were obtained from Acros Organics (Geel, Belgium). Ethanol (96% vol.) and n-dodecane (≥99%) were purchased from VWR International (Fontenay-sous-Bois, France). Prior to its use, dodecane was passed through a Supelclean™ LC-Si solid-phase extraction (SPE) column (Product no. 505374; Sigma-Aldrich, Taufkirchen, Germany) to remove impurities.

### Microalgal cultures and growth conditions

Four strains of the green model microalga *Chlamydomonas reinhardtii* that had been previously engineered to produce the sesquiterpenoid patchoulol (Lauersen et al., 2016), the diterpenoids taxadiene, casbene, or 13*R* (+) manoyl oxide (Lauersen et al., 2018), as well as their parental strain (‘UVM4’ hereafter termed wild type) (Neupert et al., 2009), were used in this work to investigate the terpenoid-extraction potential of perfluoro liquids from two-phase solvent-medium living microbial cultures. All algal cultures were maintained routinely on Tris acetate phosphate (TAP) (Gorman and Levine, 1965) agar plates under ∼50 μmol m^-2^ s^-1^ light intensity before transfer into 24-well plates containing 1 mL TAP medium, and shaken at 190 rpm on a 12h:12h dark: light (∼150 μmol m^-2^ s^-1^) cycle. After 2 d, an additional 500 μL of medium was added to each well to replenish nutrients. At 4 d, 400 μL of each culture was inoculated in 40 mL TAP in Erlenmeyer flasks at 120 rpm under the light conditions indicated above and used for subsequent experiments.

### Culture growth measurements

Culture growth was measured by recording cell densities using flow-cytometry with an Invitrogen Attune NxT flow cytometer (Thermo Fisher Scientific, UK) equipped with a Cytkick microtiter plate autosampler unit. Prior to analysis, each biological triplicate sample was diluted 1:100 with 0.9% NaCl solution. Of each diluted sample, 250 μL was measured in technical duplicates (n=2) using a 96-well microtiter plate loaded into the autosampler. Samples were mixed three times each immediately before analysis, and the first 25 μL of sample was discarded to ensure stable cell flow rate during measurement. Data acquisition stopped when 50 μL from each well was analyzed. All post-acquisition analyses and population clustering were performed using Attune NxT Software v3.2.1 (Life Technologies, USA).

Algal culture health in contact with dodecane or perfluoro liquids was measured by the variable chlorophyll fluorescence of photosystem II (PSII) with a Pulse Amplitude Modulation (PAM) fluorometer (Mini-PAM-II; Heinz Walz GmbH, Germany). Before each measurement, microtiter plates containing cultures were dark-adapted at room temperature for 15 min to ensure all PSII reaction centers were open. For each sample, the signal amplitude was adjusted before one single-turnover measurement was recorded per sample. The maximum photochemical efficiency (*F*_v_/*F*_m_) was determined using the following relationship (Eqn 1), where *F*_m_ and *F*_0_ are the maximal and minimal PSII fluorescence of dark-acclimated *C. reinhardtii* cells, respectively (Baker, 2008; Genty et al., 1989):

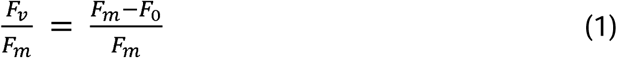

### Terpenoid capture from microalgal cultures

Two-phase living extractions were performed with dodecane, FC-40, FC-3283, and FC-770 by gentle pipetting 500 μL of each compound onto 4.5 mL liquid cultures of *C. reinhardtii* immediately after inoculation. Dodecane instantly formed an upper ‘overlay’ phase as reported in several previous studies (Beekwilder et al., 2014; Lauersen et al., 2016), while FC compounds sank under cultures to form ‘underlays’. FC compounds, however, resembled large bubbles under growing cultures and did not spread as evenly as dodecane on the surface, likely due to the liquid dynamics of water pressing on top of the FC liquids. The two-phase cultures were grown for 10 d in triplicates in 6-well microtiter plates on laboratory shakers as described above.

### Liquid-liquid extractions

To test whether the extracted patchoulol could be readily isolated from FC-3283 and enable fluorocarbon recycling, we performed liquid-liquid extractions with ethanol. Specifically, after the terpenoid capture experiments, the FC-3283 underlay from each microtiter well (∼500 μL) was recovered and transferred into a separate 1.5 mL microcentrifuge tube. The tubes were centrifuged at 3,500 x g for 5 min to separate the FC-3283 from algal cell debris. From each tube, 250 μL of the clean FC-3283 fraction was pipetted into a clean 2 mL microcentrifuge tube to which 250 μL of ethanol (96% vol.) was added. The fluorocarbon/ethanol mix was shaken at room temperature for 16 h with 1000 rpm using an Eppendorf ThermoMixer C (Eppendorf AG, Germany) to perform the liquid-liquid extraction. After the extraction, each sample was centrifuged at 3,500 x g for 5 min to separate the two liquid phases. From each sample, 90 μL of both fractions were pipetted into separate gas chromatography (GC) vials and stored at −20 °C until further analysis as described below.

### Gas chromatography

Dodecane and perfluoro liquids fractions were analyzed using a gas chromatograph equipped with a mass spectrometer and a flame ionization detector (GC-MS-FID) with minor modifications to a previously described protocol (Lauersen et al., 2016). Briefly, the GC-MS-FID analyses were performed using an Agilent 7890A gas chromatograph connected to a 5975C inert MSD with triple-axis detector. The system was equipped with a DB-5MS column (30 m × 0.25 mm i.d., 0.25 μm film thickness) (Agilent J&W, USA). The temperature profile was set to: injector (250 °C), interface (250 °C), and ion source (220 °C). 1 µL of sample was injected in splitless mode with an autosampler (Model G4513A, Agilent). Column flow was kept constant at 1mL min^-1^ with helium as a carrier gas. The initial GC oven temperature of 80 °C was held for 1 min, then raised to 120 °C at a rate of 10 °C min^-1^, followed by 3 °C min^-1^ to 160 °C, and further to 240 °C at 10 °C min^-1^, which was held for 3 min. Mass spectra were recorded after a 13.2 min solvent delay using a scanning range of 50– 750 m/z at 20 scans s^-1^.

Gas chromatograms were evaluated with MassHunter Workstation software version B.08.00 (Agilent Technologies, USA). The NIST library (National Institute of Standards and Technology, Gaithersburg, MD, USA) was used to identify patchoulol, along with verification using a purified patchoulol standard (Item: 18450, Cayman Chemical Company, MI, USA). Standard calibration curves in the range of 1–200 μM patchoulol in dodecane and FC-3283 were used for quantification. In dodecane samples only, 250 μM of *α*-humulene was additionally applied as internal standard. Extracted-ion chromatograms (XIC) with mass ranges of 91.00, 138.50, and 223.00 were used for samples with internal standard, and 138.00 and 222.00 for samples without internal standards. All GC-MS-FID measurements were performed in duplicate, and chromatograms were manually reviewed for quality control.

## Results and Discussion

Microbial engineering requires design-build-test-learn cycles as an iterative process of genetic engineering steps to identify optimal combinations of elements to drive cellular flux towards a desired product. These iterations require lengthy transformation and phenotypic screening that can be resource-heavy and laborious. Our engineering efforts in *C. reinhardtii* have relied on the use of dodecane for solvent-culture two-phase extraction of these products, which has served as a bio-compatible compound that can be readily separated and used directly in chromatography (Lauersen, 2019). Recently, it was shown that individual cells or groups of cells can be handled through encapsulation in microfluidic droplets to enable phenotyping without the need for selection agents (Yu et al., 2021). We were surprised to note that the perfluoro liquids used to make these droplets form a dense under-layer to the culture medium in large quantities. When incubated with our engineered *C. reinhardtii* cultures, we observed the accumulation of heterologous isoprenoids in these perfluorocarbons. Here, we report on and characterize the suitability of perfluoro liquids (FCs) as alternatives to currently used petroleum solvents for microbial cell isoprenoid milking.

### Potential of FC-3283 to extract terpenoids from growing microbial cultures

We first cultivated a *C. reinhardtii* strain previously engineered to produce the heterologous sesquiterpenoid alcohol patchoulol (Lauersen et al., 2016) with either a dodecane overlay or a commercially available perfluorocarbon liquid FC-3283 underlay (Fig. 1). The suitability of FC-3283 for patchoulol extraction for this culture is evident in the clear appearance of the product peak in GC-FID chromatograms in both solvents (Fig. 1a). Patchoulol was confirmed as the appropriate product by mass fractions. FC-3283 is more dense than water and was found underneath the culture medium in contrast to the lighter dodecane, which floats on top (Fig. 1b). Both solvents enabled the algal culture to grow, with cell densities increasing throughout cultivation in standard conditions and consistent linear electron flow in both, indicating healthy cultures (Fig. 1c,d). Healthy green culture is also noticeable in Fig. 1b. We then checked the suitability of FC-3283 for extraction of other heterologous isoprenoid products using strains previously engineered to product taxadiene, casbene, and 13*R*(+) manoyl oxide (Lauersen et al., 2018). Each strain generated unique products in chromatograms (Fig. 1e), as previously observed for living extraction with dodecane (Lauersen et al., 2018). The results indicated that FC-3283 is bio-compatible and amenable to accumulating heterologous isoprenoids from our engineered algal cells.

**Figure 1.**
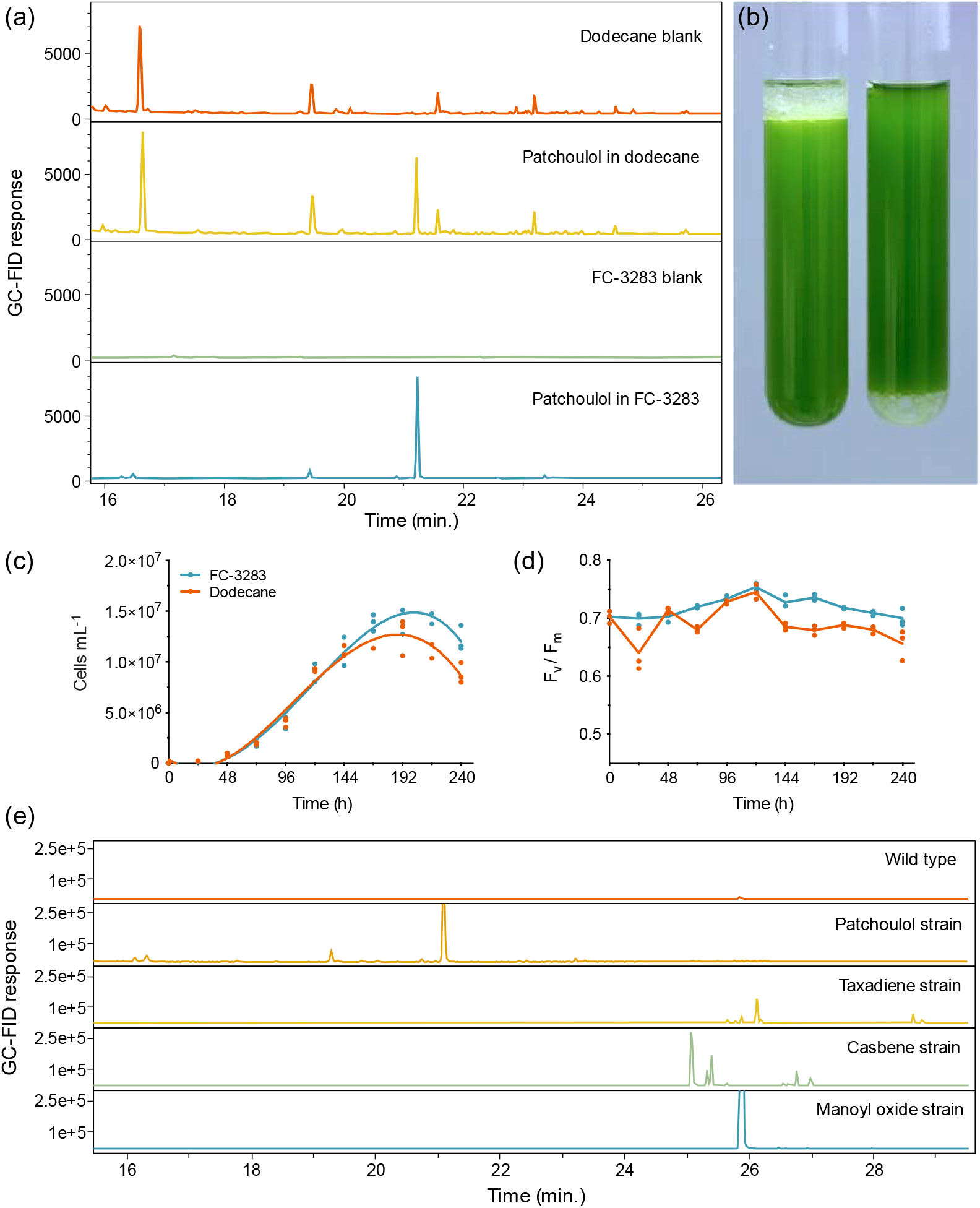
(a) Representative GC-FID chromatograms of dodecane and FC-3283 samples with and without patchoulol accumulated from incubation with growing engineered algal strains. (b) *C. reinhardtii* cultures with either dodecane overlay (left) or FC-3283 underlay (right) after 6 d on a rotary shaker with second phase. (c) Cell densities and (d) F_v_/F_m_ of *C. reinhardtii* cultures during a 10 d microtiter plate experiment with either a 5% (v/v) dodecane overlay (red line) or FC-3283 underlay (blue line) added to the culture (n=3). (e) GC-FID chromatograms of FC-3283 underlays from a *C. reinhardtii* wild type culture and various terpenoid-producing engineered *C. reinhardtii* strains harvested after 6 d cultivation.

### New options for two-phase cultivation with underlays

As a lower phase to growing liquid microbial culture, FCs present a new avenue for bio-process designs aimed at milking heterologous products. We investigated whether the lower phase could be used as a support for microbes to grow while concomitantly enabling extraction from the bio-film. We compared a shaken culture to a static culture for their abilities to produce patchoulol (Fig. 2). Both cultures grew, with the non-shaken culture accumulating a layer of cells at the medium-FC interface. Although overall patchoulol titers were considerably lower in the static (202 ± 5 μg L^-1^ culture; mean ± SD) than in the mixed culture (759 ± 73 μg L^-1^ culture) after eight days, patchoulol was found in the FC-3283 phase of both treatments already after four days of cultivation (static: 177 ± 25 μg L^-1^ culture; shaken: 263 ± 14 μg L^-1^ culture) (Fig. 2b). It seems logical that static cultures would have lower overall production rates, as only the cells close to the interface would interact with the FC. Future concepts may involve some amount of mixing and settling, or agitation of the FC liquid below the culture to increase this interaction. The ability to grow organisms at the solvent interface could enable the cultivation of mat-forming organisms and the extraction of their natural products. For example, the members of the genus *Botryococcus*, often found on the surface of stagnant waters, are slow-growing algae that can secrete alkadienes and alkatrienes or botryococene isoprenoids (Eroglu and Melis, 2010; Lozoya-Gloria et al., 2019; Templier et al., 1991). Cultures set up with larger surface area interaction between medium and FC may enable interesting process designs for milking secreted products.

**Figure 2.**
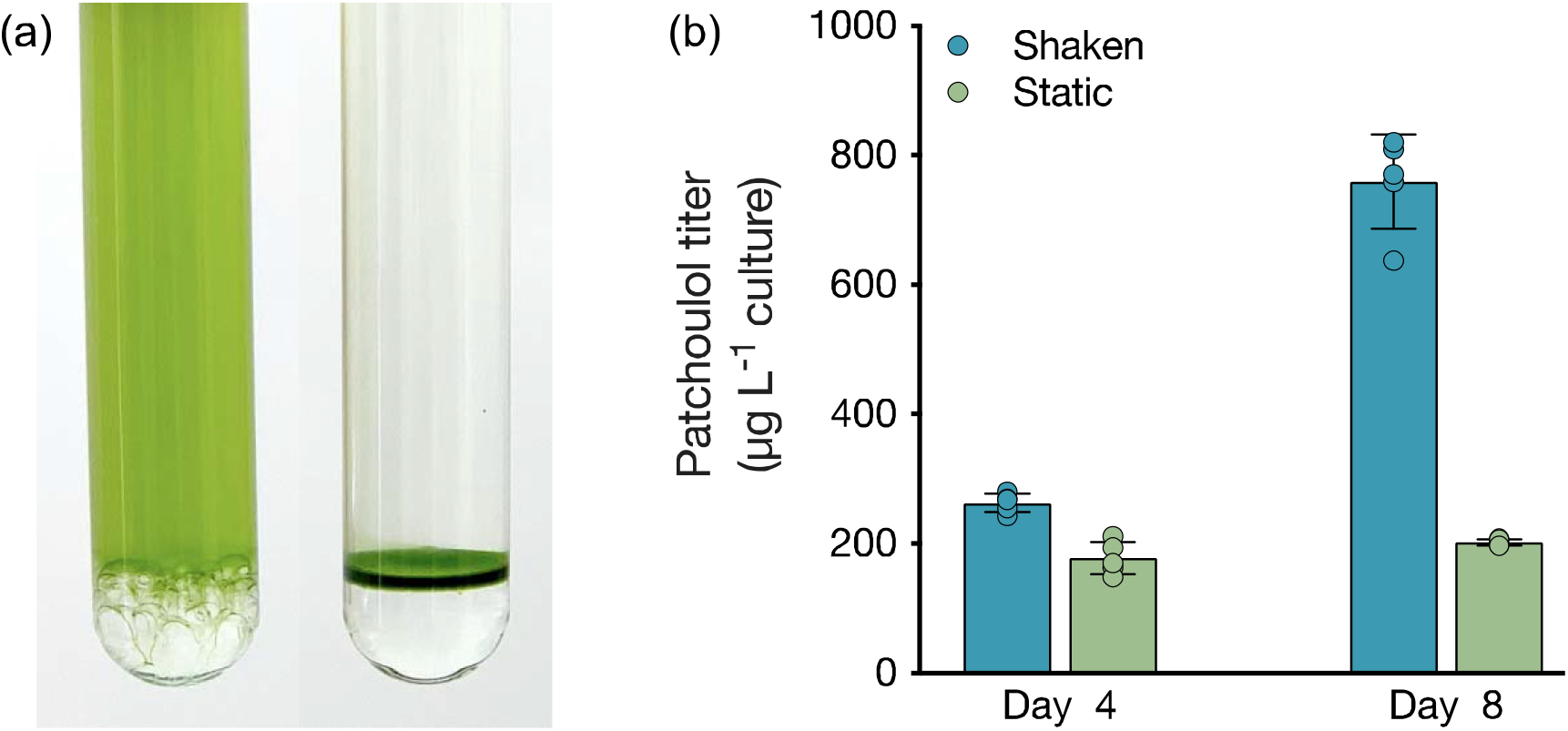
(a) Shaken (left) and static (right) *C. reinhardtii* cultures with FC-3283 underlay. (b) Patchoulol titer (μg patchoulol L^-1^ culture) in FC-3283 obtained from shaken (blue) and static (green) *C. reinhardtii* cultures at 4 d and 8 d. Values are means ± SD (n=5).

### Liquid-liquid extraction of isoprenoids from FC-3283

Once accumulated in a solvent, it is desired to extract and isolate the target chemicals and recycle the solvent for further use. With dodecane, silica-based solid phase extraction (SPE) allows patchoulol to be isolated from the solvent due to its hydroxyl group, however, separation is more challenging with non-functionalized isoprenoids. This challenge is different for every target compound, but many sesqui- and diterpenoids have similar chemical properties to the C12 dodecane, making processes like distillation or molecular separation more challenging. We sought to determine how readily we could isolate target isoprenoids from FC-3283 while regenerating the FC (Fig. 3). As it is immiscible with ethanol (Fig. 3a), we performed liquid-liquid extraction on FC-3283 which had accumulated patchoulol from a *C. reinhardtii* cultivation. Surprisingly, patchoulol partitioned readily into the ethanol phase (Fig. 3b), which phase-separated from FC3283 after extraction. We found that the process can be completed in as fast as 2-hours of incubation (data not shown), indicating the potential to obtain formulated isoprenoid product from microbial cultures in as little as two downstream process steps.

**Figure 3.**
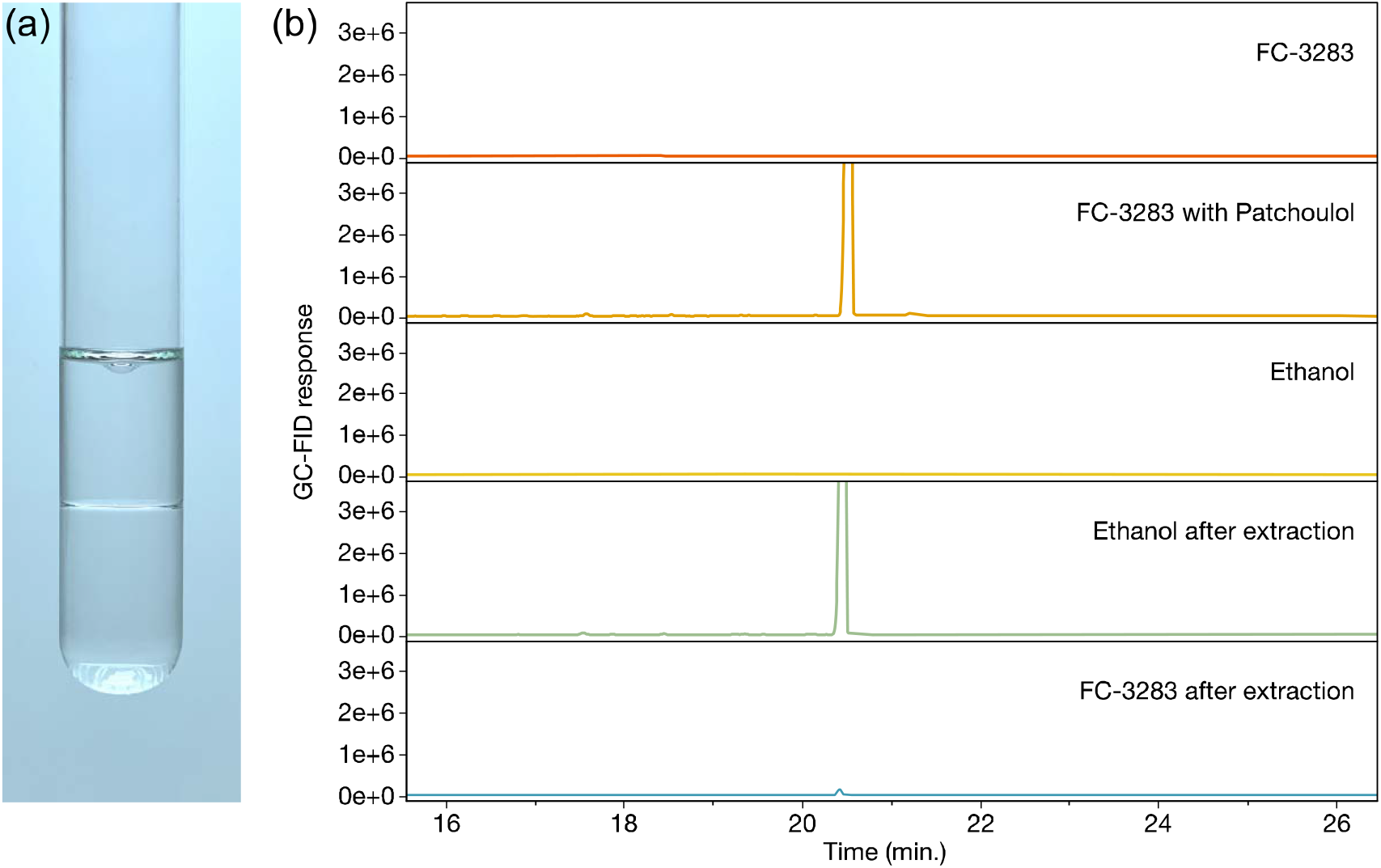
(a) Liquid-liquid extraction of FC-3283 (bottom layer) with 96% ethanol (upper layer). (b) Representative GC-FID chromatograms of liquid-liquid patchoulol extraction samples, with a clear patchoulol peak (20.4 mins) appearing in FC-3283 before extraction and in ethanol after.

### Extraction potential of other fluorinated liquids

Encouraged by the results of FC-3283 microbial milking, we cultivated *C. reinhardtii* strains producing multiple heterologous isoprenoids with other perfluorocarbon heat-transfer liquids FC-40 and FC-770 (Fig. 4). Each FC has unique physical properties and exhibited variable performance in extracting different isoprenoids from the algal cells (Fig. 4). Each fluorocarbon resulted in varying extraction rates for each isoprenoid (Fig. 4), and these behaviors will enable tailoring of bio-processes to specific product chemistries giving the operator several choices in two-phase synthetic solvents. FC-770, for example, routinely exhibited product peaks in GC chromatograms which were absent or in reduced quantities for the other FCs (Suppl. Fig. 1). Perhaps this represents a more accurate profile of side-product isoprenoids produced by these strains. Indeed, chromatograms of FCs contain much fewer background peaks and contaminants compared to n-dodecane samples which also makes characterization more straightforward (Fig. 1a). We also found that the liquid-liquid extraction with ethanol was highly effective for all analyzed isoprenoids, regardless of the FC compound used or the chemical nature of the isoprenoid, indicating a greater practical value than traditional solvents (Suppl. Fig. 2).

**Figure 4.**
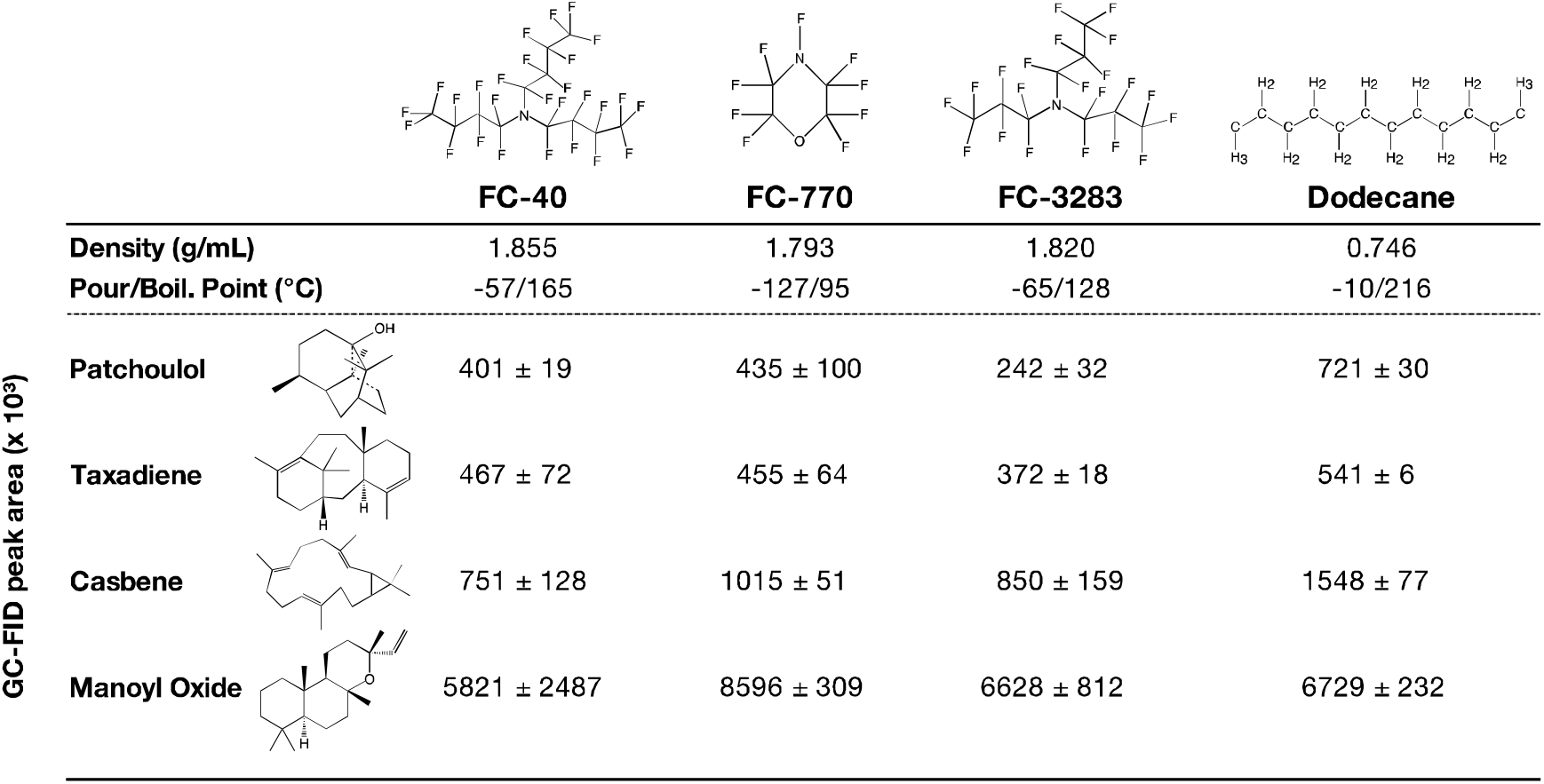
Chemical structures and properties of commercial FC liquids and dodecane that were investigated in this work. Indicated underneath are GC-FID peak areas of different terpenoids after 10 d of 2-phase living *C. reinhardtii* extraction cultures with each solvent. Peak areas represent mean ± SD (n=5). The chemical structures of FC liquids were obtained from published literature (FC-40 from Hasi et al. (2008); FC-770 & FC-3283 from Hasi et al. (2013)). All Chemical structures were drawn using ChemDraw v20.1.

Compared to dodecane, most FCs were less efficient in isoprenoid extraction from the algal cells in batch culture (Fig. 4). This can likely be an acceptable compromise in bio-process designs when considering the efficiencies of liquid-liquid extractability of the isoprenoid products from the FCs (Fig. 3). Cultures with FC-3283 overlays also exhibited higher linear electron flow rates than those with dodecane throughout batch cultivation (Fig. 1d), indicating FC-3283 is likely less aggressive as a two-phase culture system than dodecane. It could be speculated that higher efficiencies of extraction with dodecane are due to microbial lysis rather than efficient partitioning, although this aspect was not investigated here.

## Conclusions

Here, we have presented evidence that commercially available heat-transfer perfluorocarbons can be used as bio-compatible liquids for microbial cell milking of heterologous isoprenoids. The FCs tested here showed capacities for easier handling, consistent product accumulation, and ease of product isolation better than petroleum-based solvent counterparts. As dense liquids, FCs form an underlay with microbial cultures, which opens new avenues for bio-process designs, especially for slow-growing or mat-forming microbes. The use of FC liquids in microbial metabolite milking permits inert, room temperature extraction conditions and likely provides a route to well-preserved natural product extraction from microbial cells. Here, we have used engineered microalgae as a model organism, however, these results could have applicability in the synthetic biology and metabolic engineering fields for a broad spectrum of microbes.

## Author Contributions

SO performed the experiments, and contributed to the research planning and design. The research was conducted in the laboratory of KJL, who conceived the study and contributed to experiment planning. Both authors contributed to the writing of this manuscript.

## Acknowledgements

We would like to express special thanks to Najeh Kharbatia of the KAUST Analytical Core Labs for helpful early discussions, Chandrasekaren Lakshmipathy, Abdulkhalik Khalifa, and Abdullah Alabdullatif of KAUST Lab Equipment Maintenance (LEM) team for assistance in upgrading and initializing the GC-FID-MS unit. The authors acknowledge Prof. Dr. Ralph Bock and Dr. Juliane Neupert for providing *C. reinhardtii* UVM4, obtained under material transfer agreement between KAUST and the Max-Planck-Institut für Molekulare Pflanzenphysiologie Potsdam. The research reported in this publication was supported by the KAUST Impact Acceleration Funds program (grant 4238) and KAUST baseline funding awarded to Kyle Lauersen.

## References

Baker, N.R. (2008) Chlorophyll fluorescence: a probe of photosynthesis in vivo. Annu Rev Plant Biol, 59, 89–113.

Beekwilder, J., van Houwelingen, A., Cankar, K., van Dijk, A.D.J., de Jong, R.M., Stoopen, G., Bouwmeester, H., Achkar, J., Sonke, T., Bosch, D. (2014) Valencene synthase from the heartwood of Nootka cypress (Callitropsis nootkatensis) for biotechnological production of valencene. Plant Biotechnology Journal, 12, 174–182.

Beran, F., Rahfeld, P., Luck, K., Nagel, R., Vogel, H., Wielsch, N., Irmisch, S., Ramasamy, S., Gershenzon, J., Heckel, D.G., Kollner, T.G. (2016) Novel family of terpene synthases evolved from trans-isoprenyl diphosphate synthases in a flea beetle. Proc Natl Acad Sci U S A, 113, 2922–2927.

Choi, M., Cho, K. (2000) Liquid cooling for a multichip module using Fluorinert liquid and paraffin slurry. International Journal of Heat and Mass Transfer, 43, 209–218.

Daletos, G., Katsimpouras, C., Stephanopoulos, G. (2020) Novel Strategies and Platforms for Industrial Isoprenoid Engineering. Trends Biotechnol, 38, 811–822.

Eroglu, E., Melis, A. (2010) Extracellular terpenoid hydrocarbon extraction and quantitation from the green microalgae Botryococcus braunii var. Showa. Bioresour Technol, 101, 2359–2366.

Genty, B., Briantais, J.M., Baker, N.R. (1989) The Relationship between the Quantum Yield of Photosynthetic Electron-Transport and Quenching of Chlorophyll Fluorescence. Biochimica Et Biophysica Acta, 990, 87–92.

Gershenzon, J., Dudareva, N. (2007) The function of terpene natural products in the natural world. Nat Chem Biol, 3, 408–414.

Gilg, A.B., Tittiger, C., Blomquist, G.J. (2009) Unique animal prenyltransferase with monoterpene synthase activity. Naturwissenschaften, 96, 731–735.

Gorman, D.S., Levine, R. (1965) Cytochrome f and plastocyanin: their sequence in the photosynthetic electron transport chain of Chlamydomonas reinhardi. Proceedings of the National Academy of Sciences of the United States of America, 54, 1665.

Hasi, W.L., Lu, Z.W., Gong, S., Liu, S.J., Li, Q., He, W.M. (2008) Investigation of stimulated Brillouin scattering media perfluoro-compound and perfluoropolyether with a low absorption coefficient and high power-load ability. Appl Opt, 47, 1010–1014.

Hasi, W.L.J., Wang, X.Y., Cheng, S.X., Zhong, Z.M., Qiao, Z., Zheng, Z.X., Lin, D.Y., He, W.M., Lu, Z.W. (2013) Research on the compression properties of FC-3283 and FC-770 for generating pulse of hundreds picoseconds. Laser and Particle Beams, 31, 301–305.

Heider, S.A.E., Peters-Wendisch, P., Wendisch, V.F. (2012) Carotenoid biosynthesis and overproduction in Corynebacterium glutamicum. Bmc Microbiology, 12.

Jensen, S.B., Thodberg, S., Parween, S., Moses, M.E., Hansen, C.C., Thomsen, J., Sletfjerding, M.B., Knudsen, C., Del Giudice, R., Lund, P.M., Castano, P.R., Bustamante, Y.G., Velazquez, M.N.R., Jorgensen, F.S., Pandey, A.V., Laursen, T., Moller, B.L., Hatzakis, N.S. (2021) Biased cytochrome P450-mediated metabolism via small-molecule ligands binding P450 oxidoreductase. Nat Commun, 12, 2260.

Jouhara, H., Robinson, A.J. (2010) Experimental investigation of small diameter two-phase closed thermosyphons charged with water, FC-84, FC-77 and FC-3283. Applied Thermal Engineering, 30, 201–211.

Kirby, J., Keasling, J.D. (2009) Biosynthesis of plant isoprenoids: perspectives for microbial engineering. Annu Rev Plant Biol, 60, 335–355.

Lauersen, K.J. (2019) Eukaryotic microalgae as hosts for light-driven heterologous isoprenoid production. Planta, 249, 155–180.

Lauersen, K.J., Baier, T., Wichmann, J., Wordenweber, R., Mussgnug, J.H., Hubner, W., Huser, T., Kruse, O. (2016) Efficient phototrophic production of a high-value sesquiterpenoid from the eukaryotic microalga Chlamydomonas reinhardtii. Metab Eng, 38, 331–343.

Lauersen, K.J., Wichmann, J., Baier, T., Kampranis, S.C., Pateraki, I., Moller, B.L., Kruse, O. (2018) Phototrophic production of heterologous diterpenoids and a hydroxy-functionalized derivative from Chlamydomonas reinhardtii. Metab Eng, 49, 116–127.

Li, M., Hou, F., Wu, T., Jiang, X., Li, F., Liu, H., Xian, M., Zhang, H. (2020) Recent advances of metabolic engineering strategies in natural isoprenoid production using cell factories. Nat Prod Rep, 37, 80–99.

Liu, W., Xu, X., Zhang, R., Cheng, T., Cao, Y., Li, X., Guo, J., Liu, H., Xian, M. (2016) Erratum to: Engineering Escherichia coli for high-yield geraniol production with biotransformation of geranyl acetate to geraniol under fed-batch culture. Biotechnol Biofuels, 9, 131.

Lozoya-Gloria, E., Morales-de la Cruz, X., Ozawa-Uyeda, T.A. (2019) The colonial microalgae Botryococcus braunii as biorefinery. Microalgae-Physiol. Appl.

Navale, G.R., Dharne, M.S., Shinde, S.S. (2021) Metabolic engineering and synthetic biology for isoprenoid production in Escherichia coli and Saccharomyces cerevisiae. Appl Microbiol Biotechnol, 105, 457–475.

Neupert, J., Karcher, D., Bock, R. (2009) Generation of Chlamydomonas strains that efficiently express nuclear transgenes. Plant J, 57, 1140–1150.

Peralta-Yahya, P.P., Ouellet, M., Chan, R., Mukhopadhyay, A., Keasling, J.D., Lee, T.S. (2011) Identification and microbial production of a terpene-based advanced biofuel. Nat Commun, 2, 483.

Templier, J., Largeau, C., Casadevall, E. (1991) Biosynthesis of n-alkatrienes in Botryococcus braunii. Phytochemistry, 30, 2209–2215.

Wei, G., Eberl, F., Chen, X., Zhang, C., Unsicker, S.B., Kollner, T.G., Gershenzon, J., Chen, F. (2020) Evolution of isoprenyl diphosphate synthase-like terpene synthases in fungi. Sci Rep, 10, 14944.

Wichmann, J., Lauersen, K.J., Kruse, O. (2020) Green algal hydrocarbon metabolism is an exceptional source of sustainable chemicals. Curr Opin Biotechnol, 61, 28–37.

Wlodarczyk, A., Gnanasekaran, T., Nielsen, A.Z., Zulu, N.N., Mellor, S.B., Luckner, M., Thofner, J.F.B., Olsen, C.E., Mottawie, M.S., Burow, M., Pribil, M., Feussner, I., Moller, B.L., Jensen, P.E. (2016) Metabolic engineering of light-driven cytochrome P450 dependent pathways into Synechocystis sp. PCC 6803. Metab Eng, 33, 1–11.

Wolf, S., Becker, J., Tsuge, Y., Kawaguchi, H., Kondo, A., Marienhagen, J., Bott, M., Wendisch, V.F., Wittmann, C. (2021) Advances in metabolic engineering of Corynebacterium glutamicum to produce high-value active ingredients for food, feed, human health, and well-being. Essays Biochem, 65, 197–212.

Yamada, Y., Kuzuyama, T., Komatsu, M., Shin-Ya, K., Omura, S., Cane, D.E., Ikeda, H. (2015) Terpene synthases are widely distributed in bacteria. Proc Natl Acad Sci U S A, 112, 857–862.

Yu, Z.Y., Geisler, K., Leontidou, T., Young, R.E.B., Vonlanthen, S.E., Purton, S., Abell, C., Smith, A.G. (2021) Droplet-based microfluidic screening and sorting of microalgal populations for strain engineering applications. Algal Research-Biomass Biofuels and Bioproducts, 56.

Yunus, I.S., Jones, P.R. (2018) Photosynthesis-dependent biosynthesis of medium chain-length fatty acids and alcohols. Metab Eng, 49, 59–68.

Zhan, X., Zhang, Y.H., Chen, D.F., Simonsen, H.T. (2014) Metabolic engineering of the moss Physcomitrella patens to produce the sesquiterpenoids patchoulol and alpha/beta-santalene. Front Plant Sci, 5, 636.

