## Supplementary Figures 1 & 2 for "Biocompatible fluorocarbon liquid underlays for in situ extraction of isoprenoids from microbial cultures"

**Contents of this file**

Suppl. Figures 1 & 2

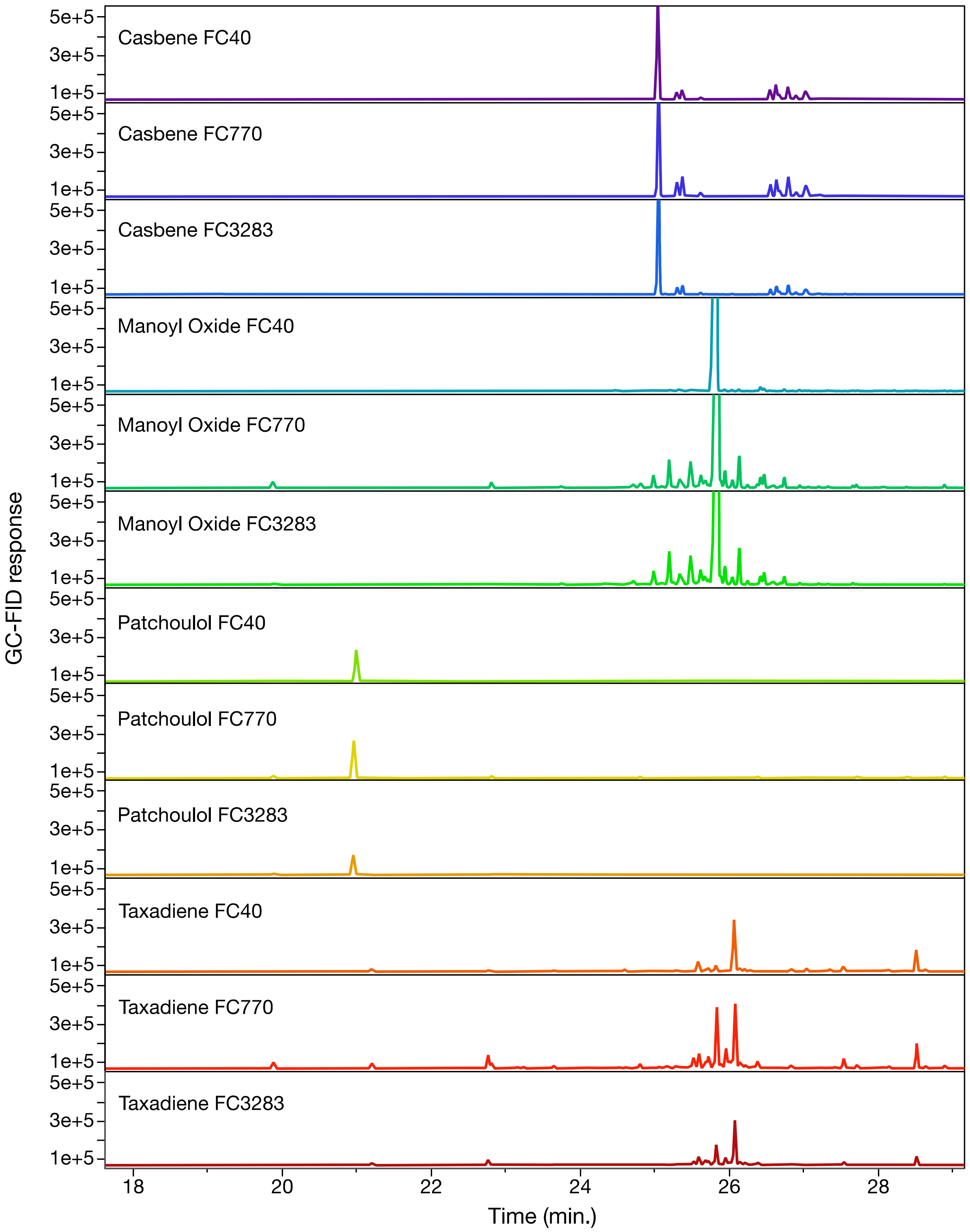


**Suppl. Figure 1.** Representative GC-FID chromatograms of FC-40, FC-70 and FC-3283 underlays after 10 d of 2-phase solvent-medium culturing with different engineered terpenoid-producing *C. reinhardtii* strains.

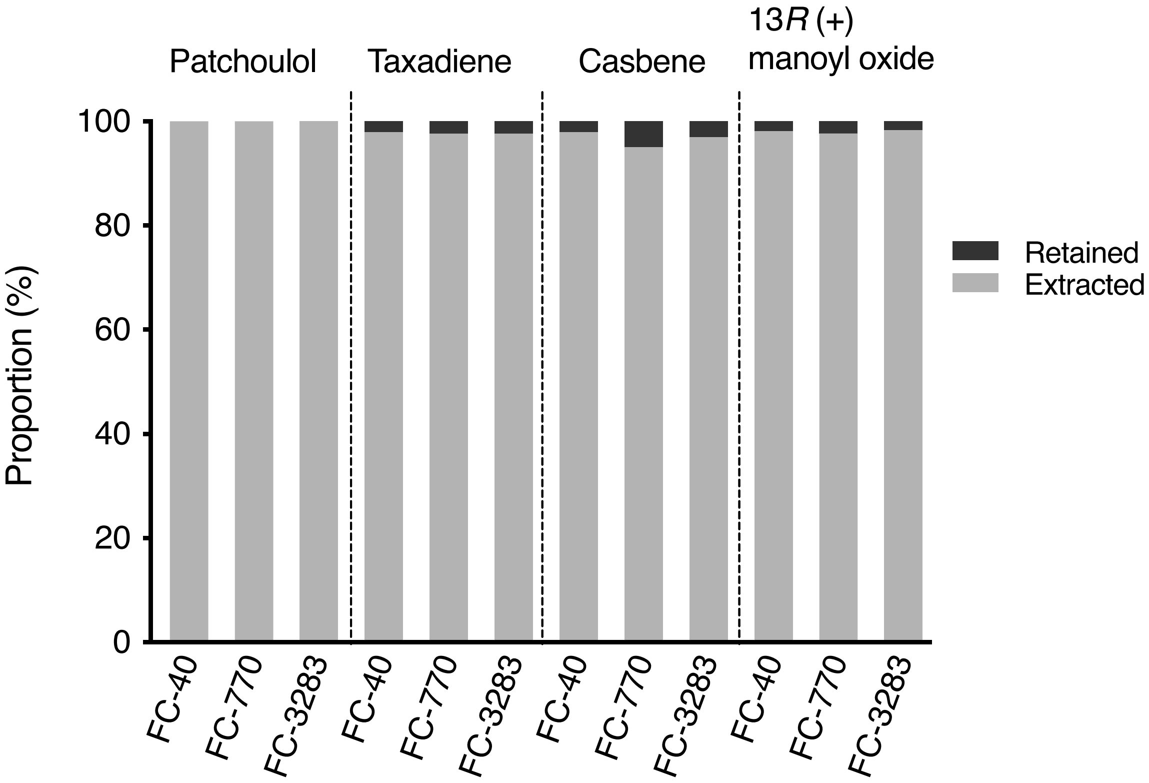


**Suppl. Figure 2.** Extraction efficiency of various terpenoids accumulated in different FCs after accumulation from two-phase engineered algal culture. Liquid-liquid extraction with 96% ethanol (1:1 v/v) for 16 h was performed on FCs. Grey bars indicate the proportion (%) of terpenoid concentration observed in ethanol after extraction compared to starting concentration in each FC.
